# Symbiont diversity and light-organ morphology in *Sepiola affinis*

**DOI:** 10.1101/2025.11.06.686942

**Authors:** Clotilde Bongrand, Raphael Lami, Marcelino T. Suzuki, Eric J. Koch

**Author notes:** <u>Communicating author</u>: Eric J. Koch. The authors declare no competing financial interests.

## Abstract

The squid-vibrio symbiosis has illuminated fundamental mechanisms of beneficial animal-microbe associations, yet the interactions within sepiolid squid in the Mediterranean Sea remain underexplored. Here we characterize the *Sepiola affinis* squid-vibrio symbiosis by combining whole-genome sequencing of light-organ isolates, confocal microscopy, and temperature-dependent growth assays. Comparative genomic analyses (ANI, phylogenomics, and functional analyses) revealed two previously undescribed *Vibrio* species associated with the *S. affinis* light organ. One species clusters more distantly from other *Vibrio* while the other species is closer to established *Vibrio* clades, with the second species exhibiting an expanded repertoire of mobile elements and Type VI secretion components, suggesting heightened capacity for genetic exchange and interbacterial interaction. Confocal microscopy of juvenile squid established that the *S. affinis* light organ comprises twelve crypts connected by pores and ducts, expanding the number of symbiotic niches relative to other sepiolid squid. In addition, fluorescently labeled isolates from the two *Vibrio* species colonized juveniles in both mono- and co-colonization patterns within crypts. Finally, growth assays across 16–24°C identified species-specific temperature differences, indicating temperature preference that may align with seasonal variability in the Mediterranean Sea. Together, these findings position *S. affinis* as a tractable model for studying how symbiont diversity, organ architecture, and interbacterial interactions contribute to the stability of a mutualistic symbiosis.

## Introduction

Animals and microbes routinely form symbiotic relationships that provide benefits for both partners [1]. These mutualisms are oftentimes integrated into aspects of host physiology such as development, immunity, and nutrition [2]. However, their inherent complexity presents a challenge for understanding the mechanisms underlying their stability. One way of addressing these challenges is through invertebrate models of symbiosis [3]. The partnership between sepiolid squid (i.e., bobtail squid) and bioluminescent *Vibrio (Aliivibrio)* bacteria (i.e., the squid-vibrio symbiosis) has served as a model for beneficial symbioses for more than 30 years [4– 6]. Primarily known for research with the Hawaiian bobtail squid *Euprymna scolopes* and its symbiont *Vibrio fischeri*, the symbiosis has yielded groundbreaking insights into how animals and bacteria initially form and subsequently maintain a mutually beneficial relationship. In addition, sepiolid squid occur throughout the world’s oceans, and many of them contain symbiotic associations with bioluminescent bacteria [7].

The most well-known symbiosis between sepiolid squid and *Vibrio* bacteria occurs within a dedicated light organ (Figure 1). When the squid first hatch, they are nonsymbiotic and acquire their bacterial symbiont from the surrounding environment [8]. The planktonic *Vibrio* migrate through sets of pores on both sides of the light organ, with each pore connecting to a duct and then ending at an epithelium-lined crypt [9]. The bacteria proliferate within the crypts, establishing symbiont populations that will remain for the lifespan of the squid. Within these crypt spaces, the host provides nutrients to the bacteria in exchange for bioluminescence that is used as counterillumination during nocturnal activity [10]. In both *E. scolopes* and *Euprymna berryi*, the bilaterally symmetrical light organ contains a total of six crypts (i.e., three on each side), while the *Sepiola robusta* light organ has two additional crypts for a total of eight [11–13].

**Figure 1.**
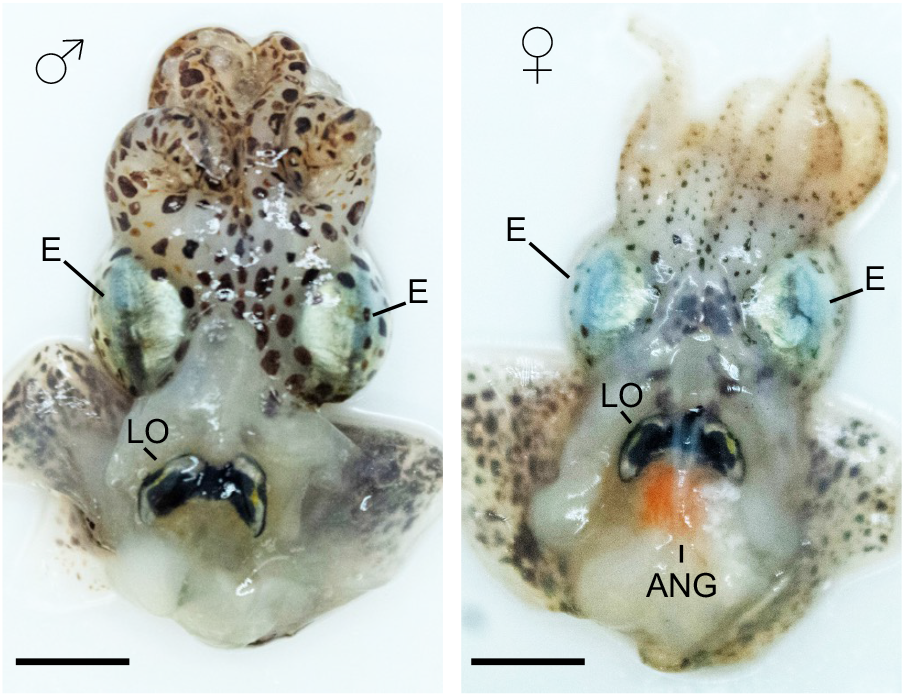
*Sepiola affinis* anatomy. Ventral dissections of a male (left) and a female (right); eyes (E) and light organ (LO) are indicated in both, and the accessory nidamental gland (ANG) is indicated in the female. Scale bars = 0.5 cm. Photographs courtesy of Alice Rodrigues.

Numerous studies have examined the diversity of the bacteria colonizing the light organ [14, 15]. In *E. scolopes*, only *V. fischeri* has been found to naturally colonize, however, there is variation at the strain level of symbionts [16]. In addition to genomic analyses, behavioral diversity among *V. fischeri* strains during colonization has also been characterized [17–19]. Furthermore, similar behaviors during colonization of *E. scolopes* are exhibited in *V. fischeri* strains isolated from other hosts, including the Japanese squid *Euprymna morsei*, suggesting conserved colonization mechanisms [20]. The light organ in *E. Berryi* can also be colonized by *V. fischeri* [13]. Interestingly, sepiolid squid from the Mediterranean Sea harbor multiple species of *Vibrio* symbionts within their light organ [21]. Specifically, both *S. robusta* and *Sepiola affinis* are colonized by *V. fischeri* and other *Vibrio spp*. [21–23]. Because of the diversity exhibited in the light organ of Mediterranean sepiolid squid, we chose to examine the squid-vibrio symbiosis within *S. affinis*.

In this study, we characterized bacterial species and examined the morphology of the *S. affinis* light organ, yielding two main findings. First, whole genome sequencing was performed on light organ symbionts, allowing for the discovery of two uncharacterized species of *Vibrio* bacteria. In addition, *S. affinis* were successfully maintained in the laboratory, allowing for the production of hatchlings and subsequent colonization with labeled bacterial strains. This allowed characterization of the *S. affinis* juvenile light organ morphology, which led to the finding that the *S. affinis* light organ contains a total of twelve symbiotic crypt spaces. These findings show the greater diversity found within the *S. affinis* light organ relative to other squid light organs and further establish it as a promising model system to study the squid-vibrio symbiosis. Continuing to explore the *S. affinis* symbiosis can bring unique insights to the evolution and ecology of mutualistic associations.

## Material and methods

### Ethics Statement

All live animal collection, husbandry, and experiments were conducted in accordance with Sorbonne Universities and the Centre National de la Recherche Scientifique (CNRS) animal welfare protocols. All maintenance, rearing, and experimental procedures with the squid adhered to the animal welfare standards of EU Directive 2010/63 on the use of animals for scientific purposes. In France, the Observatoire Océanologique has an official ethical commission agreement (A6601601) to ensure that animal welfare standards are maintained.

### S. affinis Husbandry

Adult *S. affinis* were collected at night during diving expeditions at ∼20 m of water depth near Banyuls-sur-Mer, France. The squid were maintained at the Observatoire Océanologique in 30 l aquariums with flowing natural seawater that varied with the seasonal temperatures (e.g., ∼13°C in winter to ∼24°C in summer). The animals were maintained on a 12 h:12 h light:dark lighting schedule and fed live shrimp every night. Every morning, the aquariums were inspected and any eggs laid were removed, rinsed 3 x 10 min in filtered sterilized ocean water (FSOW) and then transferred to an aquarium with 5 l of FSOW and continuous aeration. The egg clutches were housed in individual aquariums and underwent a 50% water change every day and maintained at ∼23°C. Upon hatching, the squid were removed and placed into fresh FSOW.

### Bacterial isolation

All bacterial strains used in this study were isolated from field-caught *S. affinis* collected during trips in May, June, and July of 2022. To sample the light organ symbionts, each squid was briefly rinsed 3 x 5 min in 0.22 µm FSOW, anesthetized using 2% EtOH in FSOW, and then euthanized by increasing the concentration of EtOH to 4%. The central core tissues from the light organ, which contain the symbiotic crypts, were dissected out, combined for each squid in 70% FSOW, and homogenized using a pestle to release the bacteria. Half of the bacteria were immediately mixed with glycerol diluted for a final concentration of 20% and frozen at −80°C, while the remaining homogenate underwent serial dilutions that were then grown on LBS agar [24] plates at three temperatures: 4°C, 20°C, and 25°C. Following incubation for 24 h (20°C and 25°C) or 5 d (4°C), ∼100 total colonies were selected from each light organ across the temperatures with a focus on selecting bacteria that displayed different size or color. Each colony was transferred from the agar plate to liquid LBS using a toothpick and grown at 25°C with 100 RPM shaking. After 24-48 h, cultures that exhibited visible growth had a subsample removed, to which glycerol was added to a final concentration of 20%, and were frozen at −80°C to create stocks.

### Whole genome sequencing

Genomic DNA was extracted from cultures grown in LBS during exponential phase using the DNeasy blood and tissue kit (QIAgen) according to the manufacturer’s protocol. The quantity and quality of gDNA extracted was measured using a Quantus fluorometer and a Denovix DS11 Spectrophotometer, respectively. Twelve samples (four isolates per light organ from each of three squid) were then sent to Plasmidsaurus (Eugene, OR) for whole genome sequencing using Oxford Nanopore technology, assembly, and annotation. As part of the analysis by Plasmidsaurus, all genomes underwent taxonomic analysis using Sourmash v4 and Mash [25, 26]. In addition, the Genome Taxonomy Database GTDP-tk was also used to characterize the taxonomy of each genome [27, 28]. Average nucleotide identity (ANI) analysis using FastANI v1.34 was performed against reference genomes of the top matches from Sourmash, MASH, and GTDB-tk, in addition to various other *Vibrio* strains [29]. An ANI similarity >95% was used to define the same species while >99.9% ANI similarity indicated the same strain, which were combined for further analyses [29]. Digital DNA-DNA hybridization (dDDH) was performed with the online Genome to Genone Distance Calculator [30, 31]. To construct a phylogenetic tree of the distinct strains used in the ANI analysis, Orthofinder 3.0 was used with default parameters in combination with RAxML-NG [32, 33]. After establishing the taxonomy, a Roary analysis with a 90% identity threshold was used to define genes that were shared or distinct among the newly-isolated strains [34]. To compare gene functions, eggNOG-mapper 2.1.12 was used for a COG analysis [35].

### Bacterial strains and labeling

The strains used in this study are listed in Table S1. Cultures were revived from −80°C stocks and grown overnight in LBS medium. For confocal imaging, strains were fluorescently labeled using triparental mating [36] with plasmids pVSV208 (red fluorescent protein) or pVSV102 (green fluorescent protein) [37]. Where appropriate, the medium was supplemented with chloramphenicol (2.5 µg ml^-1^) for pVSV208 or kanamycin (100 µg ml^-1^) for pVSV102.

### Colonization procedure

Bacterial strains were started from −80°C glycerol stocks onto two LBS agar plates and grown overnight at either 20°C or 25°C. Individual colonies were transferred into 5 ml of LBS and grown overnight 20°C or 25°C with 100 RPM shaking. The following morning, the culture with the highest level of growth as determined by Optical Density at 600 nm (OD_600_) was diluted 1:1000 into SWT (700 ml of FSOW, 300 ml H_2_0, 10 g NaCl, 5 g Bacto-Tryptone and 3 g yeast extract) and incubated at 23°C with 100 RPM shaking and a standard colonization procedure was followed [38]. When the bacteria had reached OD_600_ ∼0.6, the culture was serially diluted in FSOW 1:10,000 for a final concentration of ∼10,000 bacteria ml^-1^ in 150 ml of FSOW. For inoculations with two bacterial strains, each strain was first diluted to an OD_600_ ∼0.6 and then both added at concentrations of ∼5,000 CFU ml^-1^, for a total of ∼10,000 bacteria ml^-1^. Within 12 h of hatching, *S. affinis* were added to the inoculum and maintained overnight with a water temperature of ∼23°C. Following inoculation, the squid were rinsed 3 x 15 min in FSOW and then maintained in FSOW.

### Confocal microscopy

Squid were briefly anesthetized with 2% EtOH in seawater and then fixed with 4% paraformaldehyde in mPBS (50 mM sodium phosphate buffer with 0.45 M NaCl, pH 7.4) for 12 h at 4°C. After fixation, the squid were rinsed 3 x 15 min in mPBS, the mantle opened and light organ dissected out, and stained for 12 h with TOPRO-3 (Thermofisher Scientific) at a dilution of 1:1000 in mPBS and rhodamine phalloidin (Thermofisher Scientific) at a dilution of 1:40 in mPBS. Samples were rinsed 3 x 20 min at room temperature with gentle rotation and then mounted on glass slides in SlowFade Glass (Thermofisher Scientific). Imaging was performed on a Leica Sp8 Confocal Microscope within the BioPIC platform at the Observatoire Océanologique de Banyuls-sur-Mer. Post-imaging analysis and preparation for publication was performed with FIJI [39], 3D-Slicer [40], and Imaris software (Oxford Instruments).

### Temperature Growth Assays

Strains were initially grown on LBS agar, then serial dilutions were grown overnight in LBS, and the dilution at OD_600_ ∼1.0 for each strain was used further. For growth assays, the equivalent of OD=0.05 for each strain was used to inoculate 1500 ul SWT and loaded in each well of a 24-well plate. For each strain, three biological replicates (separate cultures grown on different days) were assayed, each with three technical replicates. Plates were incubated with continuous orbital shaking at 16°C, 20°C, or 24°C, and OD_600_ was recorded every 30 min for 10 h and again at 12 h and 24 h on a SpectraMax iD3 (Molecular Devices). To calculate growth rates, OD values were natural-log transformed, and exponential-phase growth rates were estimated by simple linear regression from 2–12 h. Growth rates were compared by two-way ANOVA with a Tukey’s multiple-comparisons correction in GraphPad Prism v10.6.0. An adjusted P-value < 0.05 was considered significant.

## Results

### Symbiotic Vibrio spp. from S. affinis

To examine bacterial diversity within *S. affinis*, 100 strains were isolated from each light organ of three field-caught light organs (i.e., 300 total isolates). The genomic DNA was extracted from twelve total isolates (i.e., four isolates per light organ) and sent for whole genome sequencing. The sequencing results yielded a total of eleven high-quality genomes, with one sample being excluded because of quality issues. Initial ANI analysis to compare each strain revealed that four of the genomes were nearly-identical to another sample (ANI>99.9%), and in these cases only one strain was used in this study (Figure 2A). In total, seven distinct strains were obtained from three adults: two strains from each of two adults and three strains from the third adult (Table S1). The genomes of these strains span from 4.3 to 4.6 Mb and all exhibit a GC content of 39%. The seven genomes grouped into two clusters (consisting of three and four genomes), with 98–99% ANI similarity, indicating that strains within each grouping represent the same species. Using the definition that > 95% ANI similarity represents the same species [29, 41], further analysis against *Vibrio* reference genomes confirmed that the strains are part of the genus *Vibrio* but, surprisingly revealed that none of the isolates were identified as *V. fischeri* nor *V. logei* (Figure 2A). Furthermore, none of the (Alii)*Vibrio* reference genomes exhibited ANI similarity > 95% with the *S. affinis* strains, and the most similar was *Vibrio wodanis* at ∼92% ANI similarity. However, the first grouping of 3 strains (hereafter referred to as Banyuls_Sp1) exhibited ANI similarity of ∼98% with a strain named EL58, indicating that they are from the same species. The bacterial strain EL58 was isolated in 2015 from a gorgonian coral in Faro, Portugal and identified as an *Aliivibrio sp*. [42]. The second grouping of four strains (hereafter referred to as Banyuls_Sp2) exhibited no ANI similarity >95%, with the highest similarity being with *Vibrio sifiae (92%)*. In addition to ANI analysis, dDDH reinforced the finding that the strains belonged to two species that did not share species-level similarity with other known *Vibrio* bacteria (dDDH > 70% indicate the same species) (Figure 2A, bottom). These results led to the conclusion that that the newly-isolated strains from the *S. affinis* light organ belong to two separate novel *Vibrio* species.

**Figure 2.**
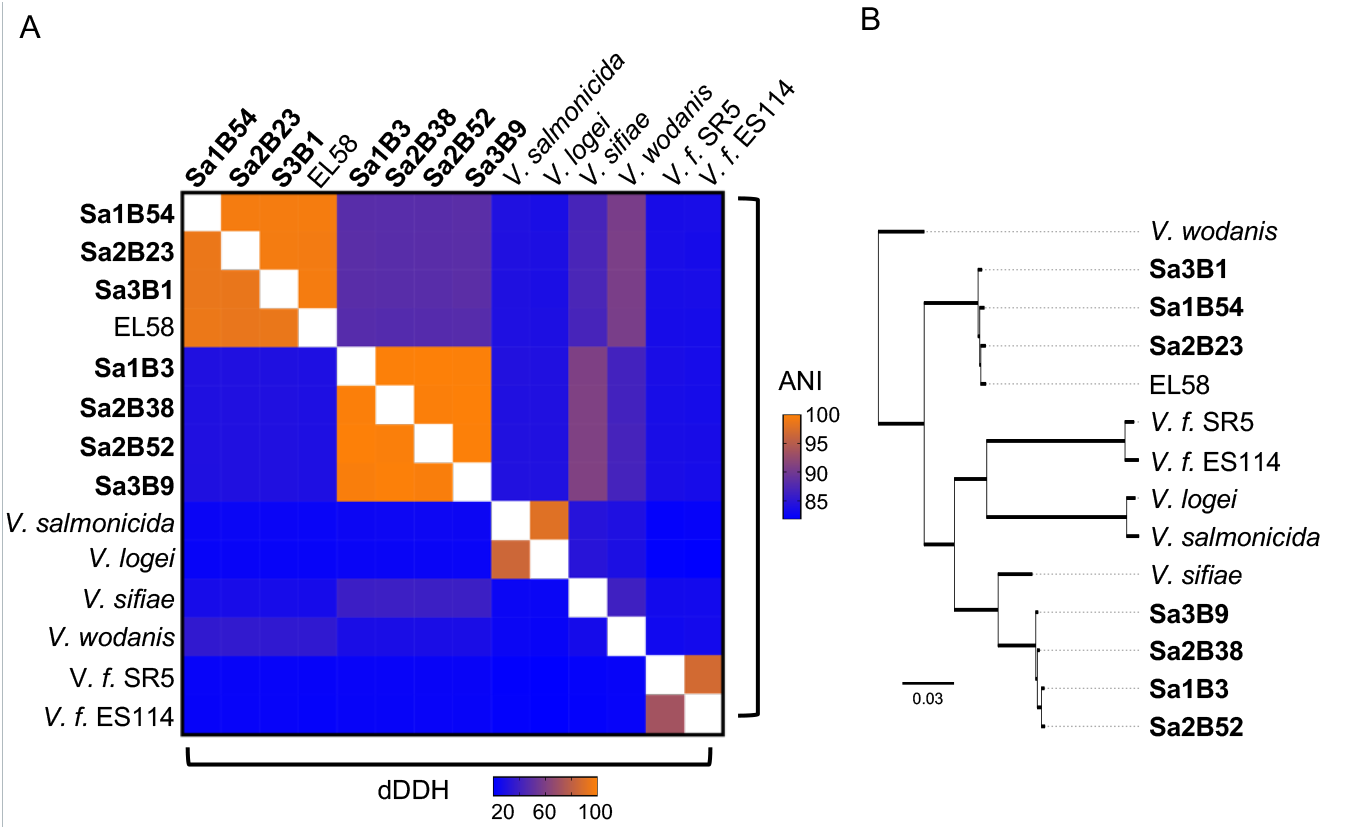
Relatedness of the *S. affinis* isolates. Strains isolated in this study are shown in bold. *V. f*. represents *Vibrio fischeri*. A) Heatmap of Average Nucleotide Identity (ANI; top) and digital DNA–DNA hybridization (dDDH; bottom) among *Vibrio* spp. strains, with colors ranging from higher similarity (orange) to lower similarity (blue). B) Maximum-likelihood phylogenetic tree inferred from 2,008 single-copy *Vibrio* genes using OrthoFinder and RAxML. The scale bar indicates the number of nucleotide substitutions per site.

To further investigate the similarity of the *S. affinis* strains relative to other *Vibrio* species, Orthofinder in combination with RAxML-NG was used to construct a phylogenetic tree from single-copy orthologs present in all of the genomes [32, 33] (Figure 2B). As expected from the ANI analysis, Banyuls_Sp1 clustered closely with EL58 and the clade separated themselves from the reference genomes, providing evidence of being from a distinct *Vibrio* species. In contrast, Banyuls_Sp2 clustered more closely to the *Vibrio* reference strains, with the closest being *Vibrio sifiae*, a bioluminescent bacterium originally isolated from surface seawater in Tokyo Bay, Japan [43]. While all strains within Banyuls_Sp2 clustered tightly together, their positioning among the reference genomes indicates that the species is more similar genomically to the other *Vibrio spp*. than Banyuls_Sp1. Overall, these results show that the bacterial strains isolated from the *S. affinis* light organ for this study represent two previously undescribed species of *Vibrio* that share a symbiotic lifestyle.

### *Comparative analyses of the Vibrio* spp

To further characterize the two *Vibrio* species, a Roary analysis was performed to compare gene presence and absence (Figure 3). Because of the ∼90% ANI similarity between the two species, we lowered the identity threshold for a shared gene from the default of 95% to 90%, as has been demonstrated in other studies that examine across different species [44]. The seven genomes contained 3913-4274 genes. From these genes, the Roary analysis built a pangenome consisting of 8107 total genes (Figure 3A). Within the pangenome, all seven strains shared 2088 genes (∼26%), which constitute the core genome. Between the two *Vibrio* species, there were 1228 (∼15%) and 1344 (∼17%) genes specific to Banyuls_Sp1 and Banyuls_Sp2, respectively. Among the individual strains, there were on average 196 to 455 (∼2%-6%) singletons or strain-specific genes.

**Figure 3.**
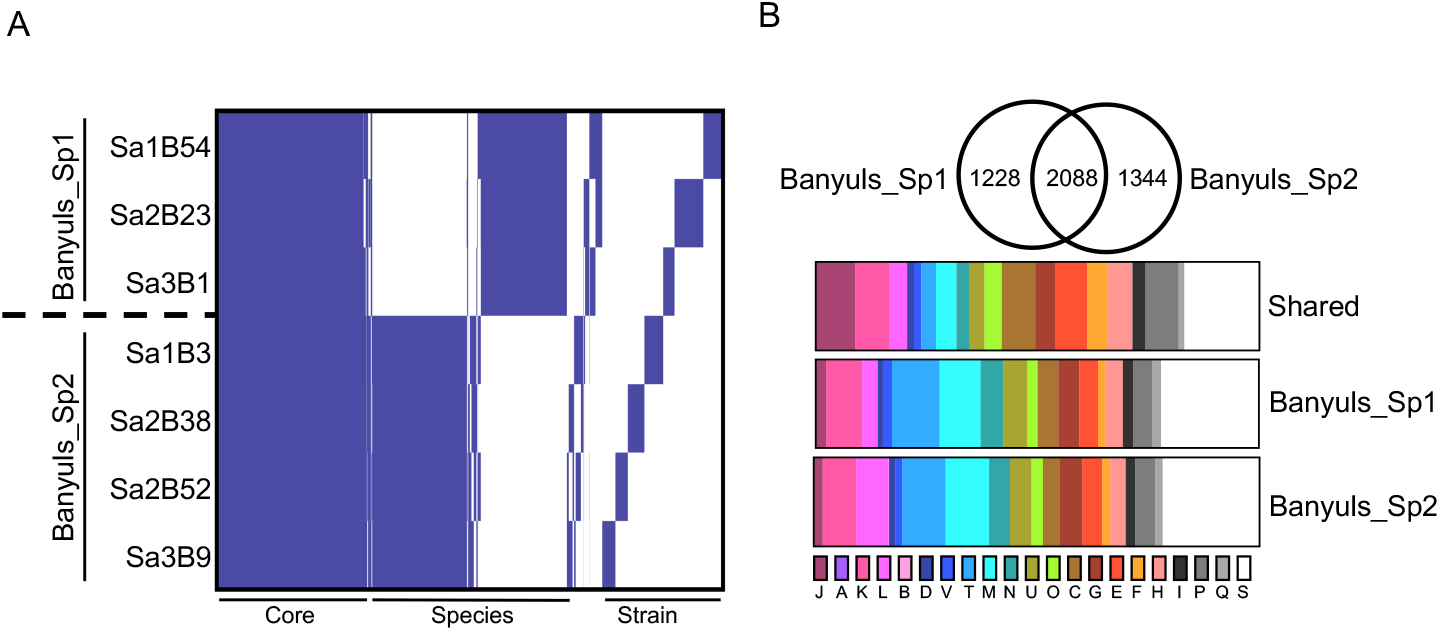
Comparison of two *Vibrio* spp. co-occurring in the *S. affinis* light organ. A) Roary presence/absence matrix for genes across the seven newly isolated symbiont strains. The two species have 2,088 shared genes (Core), ∼1,300 species-specific genes (Species), and 161–411 strain-specific genes (Strain). B) Distribution of Clusters of Orthologous Groups (COG) assignments for genes from the Roary analysis. Top: comparison of COG distributions between the two *Vibrio* spp.; bottom: distribution by COG category. COG categories: J: Translation, ribosomal structure and biogenesis; A: RNA processing and modification; K: Transcription; L: Replication, recombination and repair; B: Chromatin structure and dynamics; D: Cell cycle control, cell division, chromosome partitioning; M: Cell wall/membrane/envelope biogenesis; V: Defense mechanisms; N: Cell motility; T: Signal transduction mechanisms; U: Intracellular trafficking, secretion, and vesicular transport; O: Posttranslational modification, protein turnover, chaperones; C: Energy production and conversion; G: Carbohydrate transport and metabolism; E: Amino acid transport and metabolism; F: Nucleotide transport and metabolism; H: Coenzyme transport and metabolism; I: Lipid transport and metabolism; P: Inorganic ion transport and metabolism; Q: Secondary metabolites biosynthesis, transport and catabolism; S: Function unknown.

After establishing patterns of gene presence and absence among the two *Vibrio* species, a COG analysis by EggNOG was performed to compare functionality of the core and species-specific genes (Figure 3B). Among the core genes, the COG categories J (Translation, ribosomal structure and biogenesis), C (Energy production and conversion), E (amino acid transport and metabolism), H (Coenzyme transport and metabolism) and P (Inorganic ion transport and metabolism) are prevalent. This is expected as these categories represent basic microbial processes that are highly conserved. Among the species-specific genes, the most represented COG categories were T (Signal transduction mechanisms), M (Cell wall/membrane/envelope biogenesis), N (Cell motility) and U (Intracellular trafficking, secretion, and vesicular transport). In addition, Banyuls_Sp2 exhibited 180 genes belonging to L (Replication, recombination and repair) compared to 129 genes in Banyuls_Sp1. Among these L-category genes in Banyuls_Sp2, there were numerous genes related to transposase activity and Group II introns. The prevalence of transposase and group II intron genes indicates elevated mobile-element activity, consistent with high genetic transfer and a capacity to diversify its genome [45]. In addition, while not consistently found in the same COG category, Banyuls_Sp2 harbors a larger Type VI secretion system (T6SS) repertoire: 36 T6SS-associated genes are present in all four Banyuls_Sp2 genomes, whereas only strain Sa2B23 from Banyuls_Sp1 encodes 13 T6SS genes and the other two strains have no annotated T6SS components. These results suggest that Banyuls_Sp2 has higher potential for bacterial cell-cell communication and in DNA acquisition/exchange relative to Banyuls_Sp1.

### Influence of temperature on Vibrio spp. growth

With the knowledge that bacterial strains isolated from the light organs of *Sepiola spp*. exhibit fluctuations in growth rate according to temperature, it was next tested whether the newly-isolated *Vibrio spp*. exhibit similar patterns [21]. The growth rates for all seven strains were tested at 16°C, 20°C, and 24°C to simulate cold, medium, and warm temperatures, respectively (Figure 4). The growth rates for all strains during log-growth were compared using a two-way ANOVA and it was found that while the strains were not significantly different from one another (p-value = 0.1380), temperature was a significant factor (p-value = 0.0037). In addition, the strains belonging to Banyuls_Sp1 exhibited their highest growth rate at 16°C and lowest at 24°C, indicating that the strains grow faster at the colder temperatures (i.e., 16°C > 20°C > 24°C). Only Sa3B1 exhibited statistical significance between growth at 16°C and 24°C (adj. p-value < 0.05). For Banyuls_Sp2, the strain growth rates were highest at 20°C, but there was variability with the other temperatures, with Sa1B3 growing slowest at 16°C but the other three strains slowest at 24°C. None of these differences in growth rate were statistically significant when comparing temperatures (adj. p-value > 0.05). These results indicate that the newly-isolated *Vibrio spp*. exhibit subtle, yet consistent temperature-dependent differences in growth rate.

**Figure 4.**
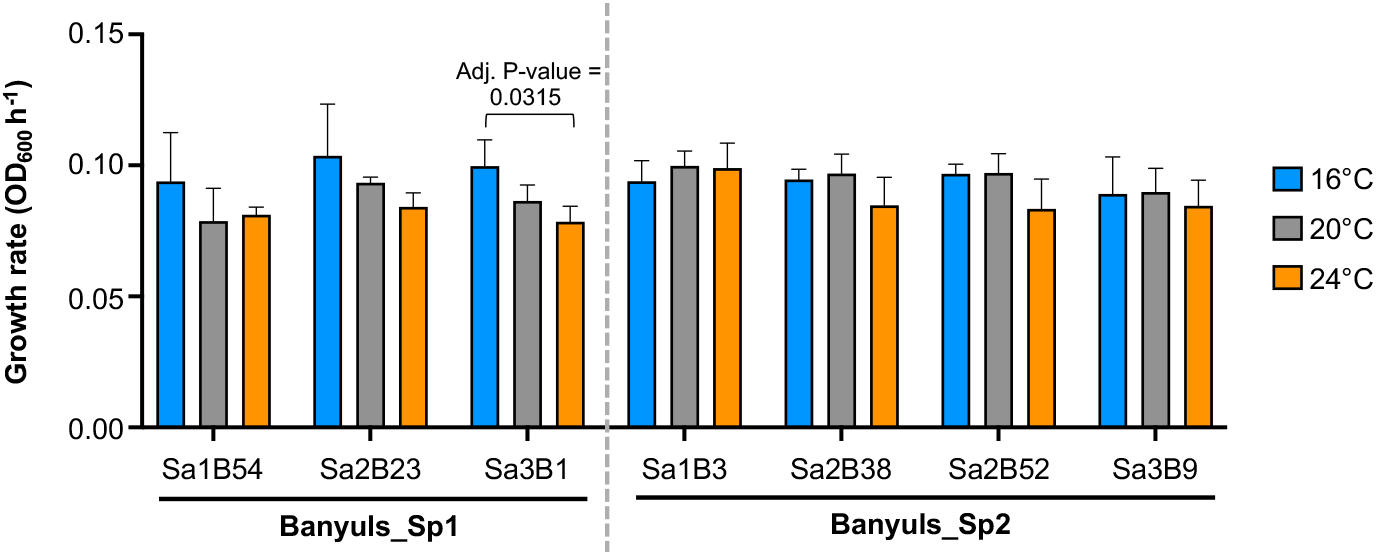
Effect of temperature on growth of *Vibrio* sp. strains. Growth rate for each strain measured at 16°C (blue), 20°C (grey), and 24°C (orange). Statistical significance of temperature effects within strains was tested by two-way ANOVA with multiple comparisons; brackets above indicate statistical significance between two growth rates. Error bars denote SD (n = 3).

### Morphology of the juvenile S. affinis light organ

The exterior appearance of the juvenile *S. affinis* light organ is consistent with other sepiolid squid [11, 12]. The *S. affinis* light organ is bilobed with a heart-shaped appearance and contains two appendages on each side: a longer anterior appendage and a shorter poster appendage (Figure 5A). At the base of each anterior appendage, a set of six pores occur that lead inside the light organ (Figure 5C). The presence of twelve pores per light organ is more than those present in *E. scolopes* and *E. berryi*, which have six pores, or *S. robusta*, which has eight [9, 12, 13]. Each pore leads to a separate duct that terminates by connecting into a crypt for a total of twelve distinct crypt spaces that will house the bacterial symbiont(s). To visualize the interior of the juvenile squid colonized by symbionts, *S. affinis* were exposed to fluorescently labelled ES114, a well-studied *V. fischeri* strain that has stability harboring fluorescent plasmids [46, 47]. Within the *S. affinis* light organ, the crypts are arranged in an overlapping formation that makes distinguishing them challenging (Figure 5B, Movie S1). The largest crypt (Crypt 1) is the most medial and closest to the ventral surface (Figure 5D). Crypts 2-5 then progressively get slightly smaller, more lateral, and more dorsal (Figure 5E-F). Crypt 6 is the smallest, most lateral (i.e., closest to the pores and appendages), and most dorsal, but was often partially or not colonized.

**Figure 5.**
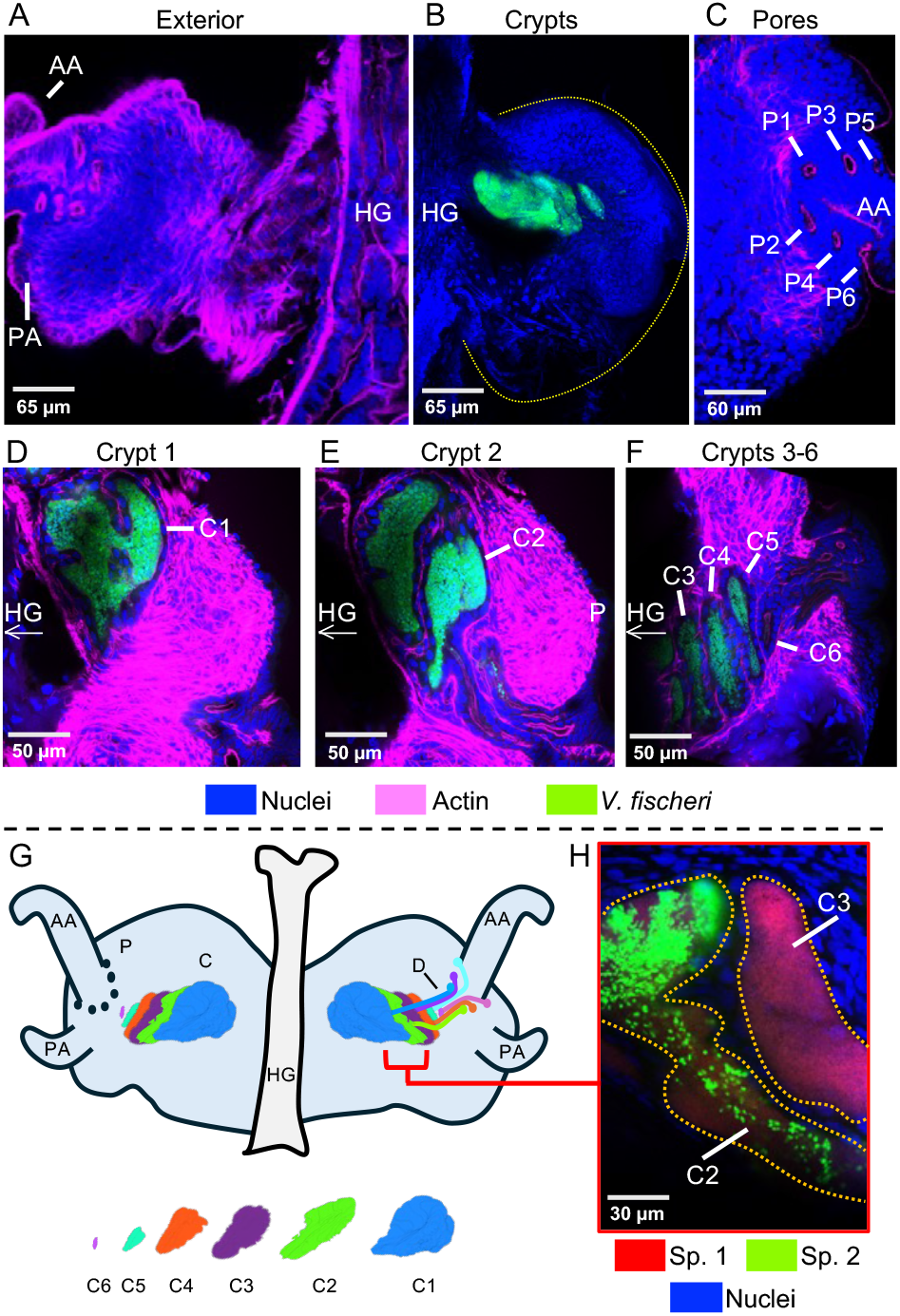
Juvenile light organ of *S. affinis* at 48 h post-colonization. Top row: (*A*) exterior view showing anterior (AA) and posterior appendages (PA) and hindgut (HG); (*B*) interior view with symbiont-containing crypts (green) and the light-organ outline (yellow dashed line); (*C*) pores (P1–P6). Middle row: (*D*) crypt 1 (C1), (*E*) crypt 2 (C2), (*F*) and crypts 3–6 (C3–C6); arrow indicates hindgut direction. Bottom row: (*G*) schematic of pores and crypts (leftside), with ducts (rightside), and individual crypts (below); (*H*) example of co-colonized (C2) and mono-colonized (C3) crypts containing newly isolated *Vibrio spp*. with Banyuls_Sp1 (Sp. 1) and Banyuls_Sp2 (Sp. 2). Scale bars, as indicated.

The six overlapping crypts create an overall architecture of near continuous symbiotic bacteria that is slightly angled outwards due to the progression from ventral to dorsal (Figure 5G). After initial characterization of the *S. affinis* light organ with ES114, some of the strains isolated from *S. affinis* were labeled with a fluorescent plasmid and tested for colonization (Figure 5F).

Through confocal microscopy, it was established that Banyuls_Sp1 and Banyuls_Sp2 strains colonize juvenile *S. affinis* when inoculated in pairs originating from the same host. Furthermore, the *Vibrio spp*. were observed both in monocolonization and co-colonization of crypt spaces. These results confirm that the newly-isolated Banyuls_Sp1 and Banyuls_Sp2 strains can colonize the *S. affinis* light organ and exhibit colonization patterns (e.g., both mono and co-colonized crypts) similar to how *V. fischeri* strains colonize *E. scolopes* [19].

## Discussion

Natural symbioses examined under laboratory conditions deepen the understanding of how mutualistic associations operate within an ecological context [48, 49]. The squid-vibrio symbiosis is a versatile model for natural animal-microbe mutualisms because it combines a host that can be maintained under both symbiotic and nonsymbiotic conditions with a bacterial partner that is highly tractable in the laboratory [50]. In addition, study of field-caught *E. scolopes* and *V. fischeri* in their native habitat of Hawaii has allowed for incorporation of ecological factors to provide a well-rounded perspective of environmentally-transmitted symbioses [15, 17]. The Mediterranean Sea harbors a wide diversity of sepiolid squid globally, providing a unique reservoir of squid-vibrio symbioses [7, 51]. Furthermore, with multiple bacterial symbionts, the squid-vibrio symbiosis in *S. affinis* provides a unique perspective into how bacterial species co-colonize and persist within the light organ. To better understand how the squid-vibrio symbiosis functions with multiple bacterial species, this study aimed to assess the microbial diversity and light organ morphology encountered within the sepiolid squid *S. affinis*.

From the *S. affinis* light organ, two species of symbiotic *Vibrio* bacteria were isolated and their genomes characterized. The discovery of these species highlights the value of whole genome approaches in bacteria, especially in *Vibrio spp*. The finding of multiple *Vibrio* species from *S. affinis* light organs was expected, as this was found in 1998 using 16S rRNA gene sequences and two species were identified as *V. fischeri* and *Vibrio logei* [21]. Subsequently, a study using a Multilocus Sequence Analysis (MLSA) approach confirmed the species identity of some *Sepiola* light-organ isolates and reclassified others [23]. Other studies have used different combinations of genes and genomes to examine the *Vibrio* taxonomy, highlighting the complexity of the genus [28, 52]. One study in particular included the genome of EL58 isolated from a Mediterranean Sea coral and grouped it with *Sepiola spp*. light organ strains, indicating that these strains represent a *Vibrio* species that exhibits a conserved role as a bacterial symbiont [52].

In addition, it is worth noting the pattern of both species being present in all three light organs examined for this study. The presence of multiple *Vibrio* species within *S. affinis and S. robusta* light organs has been linked to temperature-dependent growth patterns that were exhibited by each *Vibrio* species [21]. In contrast, *E. scolopes* is only colonized in its native habitat of Hawaii by *V. fischeri*. Therefore, the presence of two *Vibrio* species with different temperature preferences within *S. affinis* and *S. robusta* may be an adaptation to the larger seasonal temperature fluctuations in the northwestern Mediterranean Sea relative to the tropical waters surrounding Hawaii. However, the mechanisms of how the host is consistently colonized by multiple species remains unknown. Further study of the newly characterized light organ symbionts can advance our understanding of partner choice in different environments with temperature variations, providing an intriguing contrast to tropical symbioses with more constant conditions. Moreover, the *S. affinis* squid-vibrio symbiosis provides a tractable system to investigate how climate change, specifically effects related to temperature increases, could reshape marine host–microbe interactions.

The genomes of the seven *Vibrio* strains each exhibit ∼4000 genes, which is typical among *Vibrios*. Consistent within the *Vibrio* genus, strain-level diversity appears to be driven by horizontal gene transfer and gene presence/absence variation [53–55], a phenomenon also observed in other bacterial species present in horizontally acquired microbiomes [56, 57]. The functional analysis of the genomes showed that Banyuls_Sp2 contained many genes related to the exchange of genetic material (e.g., transposases and Group II intron family genes). The finding of Group II intron genes is interesting because, to the best of our knowledge, this is the first finding of the gene family in a light organ symbiont, whereas they are known to be present in pathogenic *Vibrios* such as *Vibrio cholerae* and *Vibrio paraheamolyticus* [58]. In addition, the higher prevalence of T6SS genes across all four Banyuls_Sp2 strains, relative to Banyuls_Sp1, suggests an enhanced capacity for interbacterial interaction in the former. Finally, the phylogenetic analysis provides further support for differences in genomic composition between the two species (Fig. 2B). In the phylogenetic tree, Banyuls_Sp2 clusters more closely with other *Vibrio spp*., indicating higher similarity, whereas Banyuls_Sp1 is comparatively isolated. The higher similarity of Banyuls_Sp2 to the other *Vibrio spp*. may reflect increased horizontal gene transfer (e.g., through transposases and Group II intron family genes), while Banyuls_Sp1 appears more conserved. While this study provides a first characterization of these symbiotic species, additional strains, symbiotic or planktonic, need to be collected and sequenced to better capture the diversity of the species. Furthermore, monitoring of the two species within the light organ across host maturation and seasons would yield key insights into their colonization dynamics and stability.

Finally, to visualize the symbiosis within the *S. affinis* light organ, the two *Vibrio* species were labeled with plasmids expressing a Green or Red fluorescent protein (i.e., GFP or RFP). Strains of both species were able to be successfully transformed with the plasmids, which is a reflection of the compatibility and plasticity of these *Vibrio* species. When visualizing the juvenile light organs of *S. affinis*, the general structure is similar to *E. scolopes, E. berryi*, and *S. robusta* with a bilobed organ that has ciliated appendages and a migration pathway consisting of pores leading to ducts that terminate at crypts [9, 12, 13]. However, the *S. affinis* light organ has twelve crypts, pores, and ducts, more than the six-eight present in the other squid-vibrio symbioses that have been described. The presence of nearly twice as many crypts provides more symbiotic niches for distinct populations of symbionts to grow. In *E. scolopes*, crypts are colonized by a single strain most of the time and the presence of twice as many crypts would provide opportunity for increased symbiont diversity. Thus, the greater number of crypts within the *S. affinis* light organ could be an adaptation to increase symbiont diversity and provide greater stability through seasonal fluctuations in the Mediterranean Sea. Upon examination of the newly-characterized *Vibrio* strains within the *S. affinis* light organ, both monocolonized crypts and co-colonized crypts were observed. These patterns of light organ colonization are similar to those in *E. scolopes*, where *V. fischeri* strains can co-colonize the same the same crypt or occupy different crypts [17, 18]. However, the number of light organs examined was too low for quantitative analysis. Another difference within the *S. affinis* light organ was the lack of clear antechambers in the ducts and bottlenecks at the entrance to the crypts (Movie S1), which are distinct features of both *E. scolopes* and *E. berryi*. In *E. scolopes*, the bottleneck has been linked to the daily venting behavior that occurs at dawn [59]. However, the lack of a large diel venting cycle in the *S. affinis* light organ may be the reason for the absence of the bottleneck, without a need to suddenly release the majority of the bacterial population. In the bioluminescent Japanse Pinecone Fish *Moncentris japonica*, a similar pattern of slow, continuous release of the bacterial symbionts with random larger pulses led the authors to conclude that the bacterial populations grow continuously [60]. Future studies of the native bacterial symbionts in *S. affinis* will provide evidence if there are species-dependent patterns within the light organ crypts. It is important to sample more squid and sequence more bacterial isolates to shed light on this symbiosis and be able to draw further conclusions on the microbial diversity present in *S. affinis*.

In conclusion, we deliver an in-depth characterization of the *Sepiola affinis* squid-vibrio symbiosis by examining the light-organ architecture and performing whole-genome analyses of its bacterial symbionts. Whole-genome sequencing revealed two previously undescribed *Vibrio* species within this symbiosis. Moreover, genome analyses indicate that one lineage is comparatively distinct relative to other *Vibrio spp*., whereas the other species clusters with prominent *Vibrio* clades and exhibits an expanded repertoire for genetic exchange and T6SS, implying heightened interbacterial interaction. In parallel, we characterized the juvenile *S. affinis* light-organ morphology, finding conserved migration pathways (pores→ducts→crypts) similar to other squid light organs, yet an expanded number of crypts with twelve. Taken together, these results support the hypothesis that the *S. affinis* squid-vibrio symbiosis prioritizes symbiont diversity via twelve symbiotic niches and the presence of multiple symbiotic *Vibrio* species

## Acknowledgements

This work benefited from access to the Observatoire Océanologique de Banyuls-sur-Mer, an EMBRC-France and EMBRC-ERIC. The authors are grateful to the BOSS (https://www.obs-banyuls.fr/fr/observer.html) diving and boating services (for help with animal collection) and Aquariology services (helping maintain live animals). We thank the Bio2Mar (https://bio2mar.obs-banyuls.fr) and BioPIC (https://www.obs-banyuls.fr/fr/rechercher/plateformes/biopic.html) core facilities for providing access to instrumentation and technical support. We thank Michael Fuentes of BIOM for help with maintaining live animals, and Dr. Stephanie Bertrand and Dr. Hector Escriva of BIOM for access to IMARIS software. Support was provided by Marie Skłodowska-Curie Actions under grant agreements No 101064524 to Dr. Eric Koch and No 101153902 to Dr. Clotilde Bongrand.

## Supplemental Figure Legends

**Movie S1**. Juvenile light organ of *S. affinis* at 48 h post-colonization with *V. fischeri* ES114 (green). The positioning of the hindgut (HG) and pores are indicated for orientation as well as the visible crypts (C1-C6), ducts (D1-D6), and pores (P1-P4). Counterstains are for actin (magenta) and nuclei (nuclei). Scale bar, as indicated.

**Table S1**. Strains used in the study.

