## Supplementary material for "Symbiont diversity and light-organ morphology in *Sepiola affinis*": Table S1

| Strain name | RefSeq assembly  accession number | Genome size (Mb) | Number of proteins | % GC | Host |
| --- | --- | --- | --- | --- | --- |
| *Vibrio sp.* Sa1B3 | In submission | *4.6* | *4274* | 39% | *S. affinis* |
| *Vibrio sp.* Sa1B54 | In submission | *4.5* | *4080* | 39% | *S. affinis* |
| *Vibrio sp.* Sa2B23 | In submission | *4.6* | *4220* | 39% | *S. affinis* |
| *Vibrio sp.* Sa2B38 | In submission | *4.6* | *4188* | 39% | *S. affinis* |
| *Vibrio sp.* Sa2B52 | In submission | *4.5* | *4181* | 39% | *S. affinis* |
| *Vibrio sp.* Sa3B1 | In submission | *4.3* | *3913* | 39% | *S. affinis* |
| *Vibrio sp.* Sa3B9 | In submission | *4.5* | *4113* | 39% | *S. affinis* |
| *Vibrio sp.* EL58 | GCF_900312675.1 | *4.3* | *3855* | 39% | *Eunicella labiata* (coral) |
| *V. salmonicida* LFI1238 | GCF_000196495.1 | *4.6* | *4 286* | *41%* | *Fish* |
| *V. logei* 1S159 | *GCF_001691055.1* | *4.6* | *4111* | *39%* | *N/A* |
| *V. sifiae* NBRC 105001 | *GCF_002954715.1* | *4.7* | *4175* | *38%* | *N/A* |
| *V. wodanis* | *GCF_000953695.1* | *4.6* | *4168* | 38% | Atlantic salmon |
| *V. fischeri* SR5 | *GCF_000241785.1* | *4.3* | *3810* | 38% | *S. robusta* |
| *V. fischeri* ES114 | *GCF_000011805.1* | *4.3* | *3819* | *38%* | *E. scolopes* |

**Table 1**. Strains used in the study.
